# Sparse spike trains and the limitations of rate codes underlying rapid behaviour

**DOI:** 10.1101/2022.01.28.478240

**Authors:** Joseph M. Fabian, David C. O’Carroll, Steven D. Wiederman

**Affiliations:** School of Biomedicine, The University of Adelaide; Department of Biology, Lund University

## Abstract

Animals live in highly dynamic worlds, utilising sensorimotor circuits to rapidly process information and drive behaviours. For example, dragonflies are aerial predators which react to movements of prey within tens of milliseconds. These pursuits are likely controlled by identified neurons in the dragonfly, which have well-characterized physiological responses to moving targets. Predominantly, neural activity in these circuits are interpreted in the context of a rate code, where information is conveyed by changes in the number of spikes over a given time. However, such a description of neuronal activity is difficult to achieve in real-world, real-time scenarios. Here, we contrast a neuroscientists’ post-hoc view of spiking activity with the information available to the animal in real-time. We describe how performance of a rate code is readily overestimated and outline a rate code’s significant limitations in driving rapid behaviours.

## Introduction

Sensory neural circuits take time to process and react to external stimuli. Since time is a very valuable resource, particularly for animals that live in highly dynamic environments, brains have evolved strategies to minimise processing delays. For example, dragonflies pursue their prey in acrobatic aerial pursuits, reacting to turns by prey in just 47 ms (Mischiati et al., 2015). In such a short period of time, spiking neurons can only generate a small number of impulses to inform downstream control systems. Visual neurons in the motion and target detecting pathways of flying insects have been shown to have latencies in the range of 25-40 ms, usually quantified by averaging across many stimulus presentations (Warzecha and Egelhaaf, 2000; Gonzalez-Bellido et al., 2013; Nordstrom et al., 2011; Supple et al., 2020). However neurons in the brain do not have the luxury of such averaging of technical replicates to eliminate the sparseness and stochastic variance present in spike trains. Thus, while post-hoc analysis might be able to identify rapid responses from a dataset, the real-time information which an animal has access to might be significantly less informative.

When a behaviour must be controlled on timescales too short for a single neuron to fire sufficient spikes, one solution is a population code that pools the outputs of multiple neurons. If these neurons have the same tuning properties, the mean response of one neuron across many trials can then be reasonably used to infer an estimate of the mean population response to one individual stimulus presentation (Rieke et al., 1997). However, this assumption requires evidence that a large homogenous cell population exists, which is not always the case. For example, high order neurons in the target detecting pathway of dragonflies appear to be heterogeneous in terms of receptive field location, direction tuning and spiking statistics (O’Carroll., 1993; Geurten et al., 2007; Dunbier et al., 2012; Evans et al., 2020; Fabian and Wiederman, 2021). In such a system, pooling across a population of neurons may still occur, but many repetitions from a single neuron cannot be used to infer that potential real-time population activity, and doing so overestimates the performance of rate code signals.

In this paper, we use recordings from the dragonfly visual system to evaluate the limitations of implementing an effective rate code from individual spike trains in identified neurons. We highlight differences between the view of an experimenter and an organism, noting that this conflation has caused researchers to overestimate the performance of neuronal circuits.

## Materials and Methods

We recorded intracellularly from the target selective neuron CSTMD1 (Geurten et al., 2007) in 14 immobilised wild-caught dragonflies (*Hemicordulia tau*) while presenting small moving targets (2°x2°) on a 165 hz monitor. Test targets appeared within the receptive field, and drifted at 80°/s. In primed trials, the test target trajectory was preceded by a 500 ms target trajectory terminating at our test target location, either 150 or 300 ms prior to our test target appearance. Data was sampled at 5000 hz, and responses analysed offline in MATLAB.

Examples of neural response quantification were generated from CSTMD1’s response to 41 repetitions of an identical drifting target in each recording (only 20 repetitions are displayed in raster plots for clarity). Peristimulus Time Histograms (PSTHs) were produced by binning spikes into 1 ms or 20 ms bins, before calculating spike rate. Inverse inter-spike interval (ISI) plots were produced by defining instantaneous spike rates as the reciprocal of each ISI, before resampling neural activity to a constant sample rate of 1000 hz, allowing us to average data across multiple trials. Spike probability was calculated by binning spike times into 1 ms bins, such that each bin in each trial has either 0 or 1 spikes, and calculating the probability of a spike for each 1 ms bin across all trials in the same neuron.

To define the latency of responses of CSTMD1 to the presentation of a stimulus, we adapted a noise-based threshold approach from Warzecha and Egelhaaf (2000). This approach is sensitive to the distribution of spontaneous spike rates, so we pooled data from 1008 1-second long recordings of CSTMD1’s spontaneous activity from 12 dragonflies. We determined a spike rate threshold equal to two standard deviations above the mean spontaneous rate, and define latency as the first time point where the mean response of a PSTH calculated from CSTMD1s responses to 432 target presentations in the same 12 dragonflies (1 ms bins) surpasses this value. We generate cumulative spike count plots from the same set of trials, by determining the proportion of trials which have produced a given spike count at each point in time.

To quantify the distribution of spontaneous spiking activity we performed an autocorrelation of 2500 spontaneous spikes from CSTMD1 in one dragonfly, where for every spontaneous spike, we plot the time interval of the previous, current, and next spike. We also calculate the correlation of the interval of the previous spike with the interval of the 2^nd^ through 10^th^ following spikes.

## Results

### Quantifying neural responses to stimuli

We recorded intracellular activity from CSTMD1 while presenting a series of salient moving targets. Following presentation of a target that appears in motion within the receptive field, the spiking activity increases (Fig. 1A) until the stimulus disappears. If we repeat this same stimulus many times in the same neuron, the pattern of spikes across each trial is similar but not identical (Fig 1B). We can eliminate the individual variations between trials by pooling responses across many technical replicates. The mean response of a neuron is most commonly displayed by computing a peristimulus time histogram (PSTH), where spikes from all trials are binned according to their time (20 ms bins), and the spike count in each bin is converted to instantaneous spike rates (Fig. 1C). This produces an attractive description of a neuron’s spiking activity when used on large enough datasets. An alternative approach is the Inverse-Interspike Interval plot (ISI), where the interval between each pair of spikes is used to define an instantaneous spike rate (Fig. 1D). The ISI is updated each time a new spike occurs, with the resulting data averaged across trials. A third approach is the spike probability function, where responses are defined by the probability that a neuron fired an action potential at each time point in our experiment (Fig. 1E, 1 ms bins). When averaged across enough technical replicates, this approach will converge on the PSTH, but it could be argued that it presents spiking data in units with less potential to be misleading.

**Figure 1:**
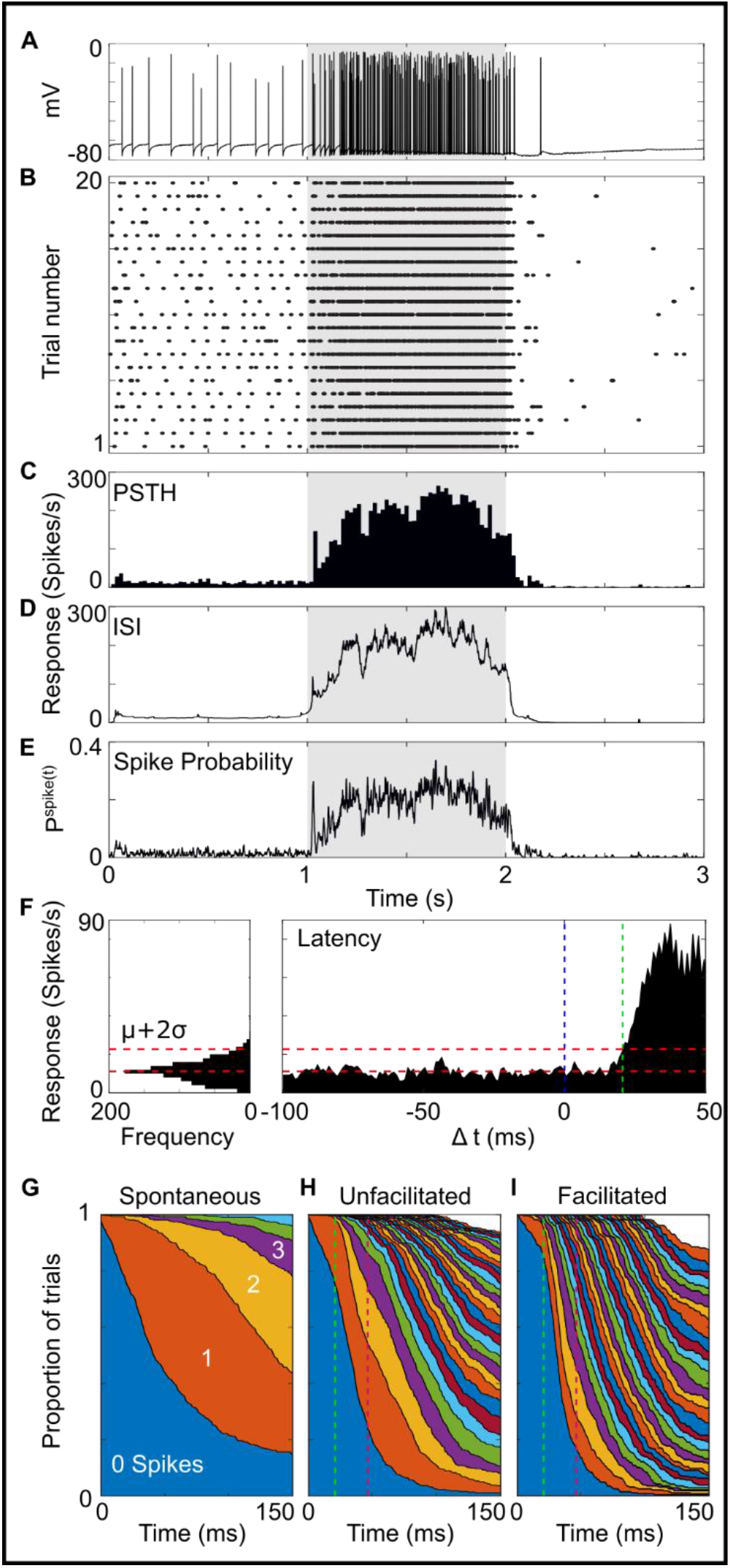
Quantifying neuronal responses in visual neurons. A) An intracellular recording from CSTMD1 during the presentation of a moving target (grey shaded region indicates peristimulus duration). B) Spike raster displaying 20 repeats of an identical stimulus from the same neuron. C-E) A PSTH (20 ms bins), an Inverse Interspike Interval plot and a spike probability function (1 ms bins) computed from the spiking data in B. F) Calculating the response latency with the approach of Warchecha and Egelhaaf (432 trials from 12 dragonflies). G-I) The same data as F, displaying the proportion of trials with a given cumulative spike count as a function of time, in the absense (G) and presence (H unprimed, I primed) of a moving target stimulus. Green and magenta dashed lines indicate the 21 ms latency from F, and the 47 ms behavioural delay from Mischiati et al (2015).

With respect to pursuits, the temporal latency between stimulus presentation and neural response is an important measure. For neurons which lack spontaneous activity this can be defined as the arrival time of the first spike. In neurons that fire spontaneously, the latency becomes a statistical problem where we must determine the earliest time when neural responses differ significantly from the pre-stimulus condition, e.g. by surpassing a threshold based on a spike distribution of spontaneous activity. We define this threshold as 2 standard deviations above the mean rate from 1008 1-second-long samples of CSTMD1’s spontaneous activity across 12 dragonflies (dashed red lines, Fig 1F). The first time where the mean response to moving targets surpass this threshold is defined as the latency (dashed green line).

All of these measures exist in the laboratory post-hoc, but none of them exist in a biological pursuit. That is, the analytical methods may be valuable for the understanding of neuronal circuits, however address a fundamentally different problem to how a neuronal circuit interprets real spike trains. To demonstrate this issue, we used the same dataset in Fig. 1F and determined the cumulative number of spikes at each time point in each trial. In the absence of a stimulus, spikes slowly accumulate over time (Fig. 1G). When a target is presented this process is accelerated (Fig. 1H), and when that target is preceded by a continuous target trajectory, predictive gain modulation (Wiederman and Fabian et al., 2017) accelerates it even further (Fig. 1I). Through post-hoc analysis of this dataset we detect an increase in spike rate after 21 ms, but at that point in time 76% of trials contained no new spikes, and only 1.4% of trials had two or more, the minimum requirement to define a real stimulus specific rate (green dashed line). Therefore in most cases, 21 ms following the presentation of a target the responses of CSTMD1 are meaningless, with respect to real-time control of behaviour. During pursuits, on average dragonflies respond to prey movements in just 47 ms (Mischiati et al., 2015, magenta dashed line). Even if we assume that all processing stages following STMD neurons are instantaneous, 47 ms following stimulus onset only 53% of trials contain two or more spikes. The event of 2 spikes in a 47 ms period also occurred in 5% of the spontaneous trials. Based on this data, if 2 spikes are insufficient to drive behaviour then most trials will result in false negatives, and the alternative where 2 spikes is sufficient, will lead to a false positive signal from spontaneous activity on average every 940 ms of a dragonfly’s life. In either case, this would not be a very desirable result.

### The problem of spontaneous activity

Many neurons, including CSTMD1, fire action potentials at low rates in the absence of a stimulus (Geurten et al., 2007). Spontaneous activity is often described as tonic, and as described in figure 1, a background noise which stimulus driven responses must surpass for detection. Researchers often calculate the mean spontaneous spike rate over large windows of time and compare this with the mean responses to a stimulus (Nordstrom et al., 2011; Wiederman and Fabian et al., 2017). However, neurons do not output spike rates, they output spikes which release neurotransmitter. Since spontaneous and stimulus driven spikes are identical at behaviourally relevant time periods, they can only be distinguished by their temporal patterns. We recorded prolonged periods of spontaneous activity in CSTMD1 (Fig. 2A) and autocorrelated spike intervals to evaluate their temporal statistics. For each spike in a train we plot the interval from the previous spike (i-1) and the next spike (i+1) (Fig. 2B). We see that spontaneous activity contains a broad distribution of spike intervals (Fig. 2C), rather than at a constant, tonic rate. To determine how rapidly spike intervals in spontaneous activity change, for 19635 spontaneous spikes, we plot the previous interval (i-1) against the next interval (i+1), the 3^rd^ next interval, and the 5^th^ next interval (Fig. 2E). Immediately neighbouring intervals are weakly correlated (R^2^ = 0.12), and this correlation quickly drops for successive spikes in a train. If spontaneous activity is not tonic and poorly correlated in time, over very short time intervals its rate can’t easily be defined, and therefore it is difficult to distinguish spontaneous and stimulus generated responses on behaviourally relevant timescales.

**Figure 2:**
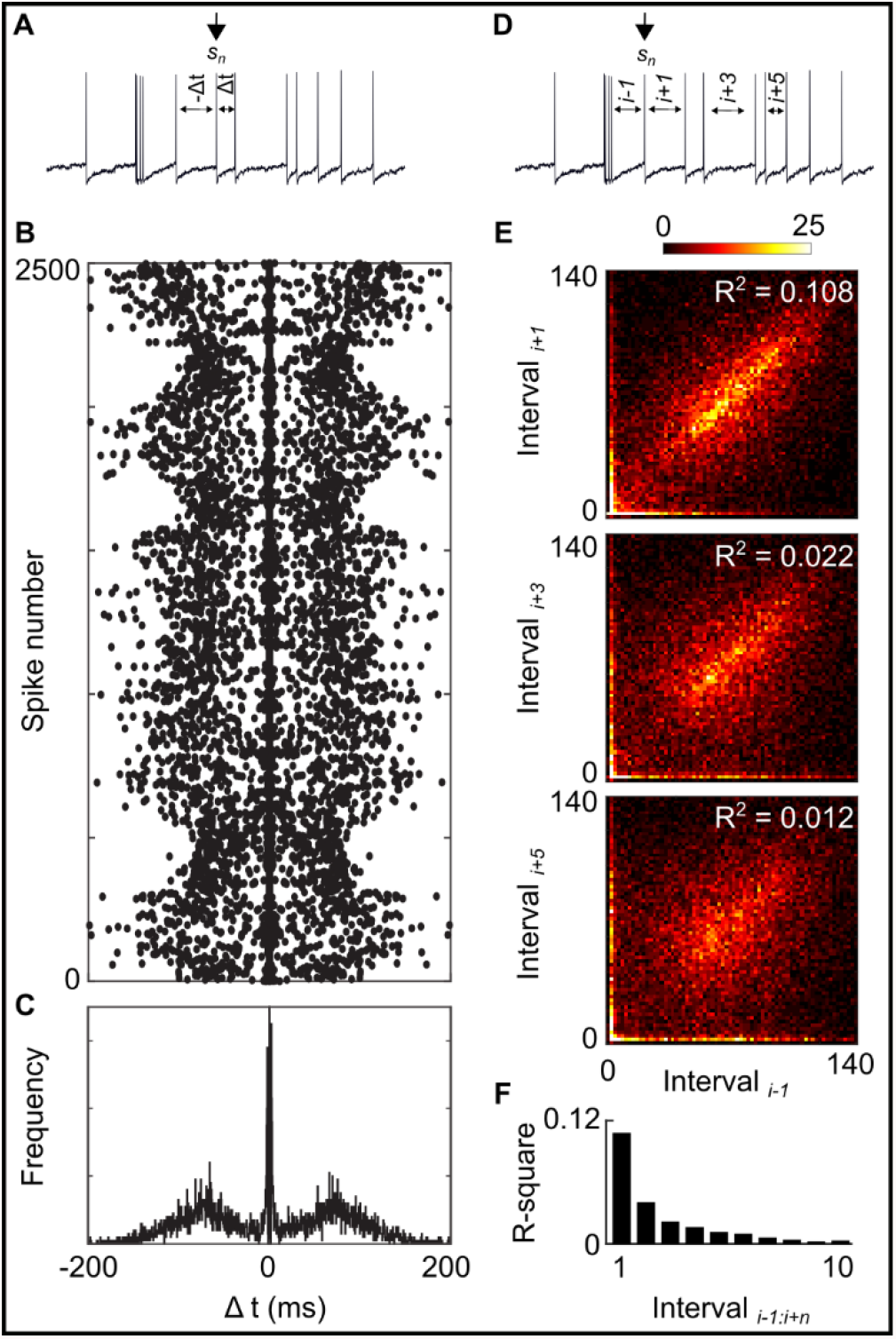
Spontaneous activity and the lack of regularity. A) A raw train of CSTMD1’s spontaneous spiking activity, defining the process of autocorrelation. B) Autocorrelation of 2500 spontaneous spikes from a single CSTMD1 recording. C) A histogram of the data in B, showing the distribution of spike intervals. D) The approach for correlating spontaneous spike intervals. E) The previous interval (i:i-1) plotted against the next interval (i:i+1), the 3^rd^ next interval, and the 5^th^ next interval for 19635 spontaneously generated spikes across 12 dragonflies. F) Spike intervals are poorly correlated, and this correlation rapidly drops further through a spike train.

## Discussion

Our detailed understanding of brain function relies on the accurate quantification of neural activity. The sparse spike trains that sensory neurons generate can be interpreted in varying frameworks, including rate or temporal codes. However, this decoding of spike trains into an informative signal is not trivial, with significant limitations on processing especially in context of sensorimotor latencies. We have demonstrated that data collected across many technical and biological replicates can overestimate the performance of neuronal circuits. Spike rates in individual trials share little resemblance with the mean PSTH, and in the short time windows where animals like a dragonfly must make decisions, simply distinguishing responses from spontaneous activity is difficult. Control theories derived from large, replicated datasets may seem enticing, but are often nonsensical in more realistic, single-trial conditions.

Answering neuroscience questions requires robust approaches for quantifying spike trains, so how should we do this? Deriving any measure from a sparse spike train transforms data in ways which could be misleading, so we need to be conscious of our approach. Spike rates are the easiest metric to calculate and the most versatile approach. Independent of the coding scheme being implemented, at a fundamental level all codes convey information by varying the timing of spikes. Thus, even if a neuron is not implementing a rate code, changes in neural signals will tend to be associated with some change in spike rate. However this analysis should be interpreted as a representation of a neuronal signal, not necessarily the true neuronal signal (see Brette, 2015). Rate codes have significant limitations, especially when temporal resolution is important, as is the case for rapid behaviours. Rate codes are sluggish because multiple spikes are required to define a rate, and spikes in a spike train can be separated by tens or even hundreds of milliseconds.

Given the limitations of rate coding, how are rapid behaviours controlled from small populations of neurons? We know that whatever solutions dragonflies use, it is remarkably effective. Spike timing codes, or bursting codes can permit faster reaction times than rate codes, because the relevant signal is contained in fewer spikes. Recent work found that when a dragonfly is presented with a salient stimulus, CSTMD1 fires spikes in a series of bursts (Fabian and Wiederman, 2021). The neurons thought to lie downstream of STMD neurons, the TSDNs, fire extremely sparse trains, with an ideal stimulus eliciting only a handful of spikes (Olberg, 1986; Gonzalez Bellido et al., 2013). These abnormal spiking patterns in the target detecting pathway suggests that pursuit is likely controlled on the basis of individual spikes across a population of neurons rather than a spike rate code, most likely via some form of temporal code. To understand the control of rapid and accurate behaviours, future work must study spike timing in detail, and focus on distinguishing practical measures of neural activity from realistic neural codes and control systems.

## Acknowledgements

This research was supported by the Australian Research Council’s Future Fellowship scheme (FF180100466) and the Swedish Research Council (VR 2018-03452). We thank the manager of the Adelaide Botanic Gardens for allowing insect collection and behavioural recordings.

## References

1. Brette, R. Philosophy of the Spike: Rate-Based vs. Spike-Based Theories of the Brain. 2015. Front. Syst. Neurosci. 9.

2. Dunbier JR, Wiederman SD, Shoemaker PA, and O’Carroll DC. Facilitation of dragonfly target-detecting neurons by slow moving features on continuous paths. 2012. Front Neural Circuits. 6:79.

3. Evans, BJE., Fabian, JM., O’Carroll, DC., and Wiederman, SD. Dragonfly visual neurons selectively attend to features in naturalistic scenes. 2020. bioRxiv.

4. Fabian, JM., Dunbier, JR., O’Carroll, DC. and Wiederman, SD.; Properties of predictive gain modulation in a dragonfly visual neuron. 2019. J. Exp. Biol. 222:jeb207316.

5. Fabian, JM., el Jundi, B., Wiederman, SD., and O’Carroll, DC. The complex optic lobe of dragonflies. 2020. bioRxiv.

6. Geurten, BRH., Nordström, K., Sprayberry, JDH., Bolzon, D M., and O’Carroll, DC. Neural mechanisms underlying target detection in a dragonfly centrifugal neuron. 2007. J Exp Biol 210, p3277–3284.

7. Gonzalez-Bellido, PT., Peng, H., Yang, J., Georgopoulos, AP., and Olberg, RM. Eight pairs of descending visual neurons in the dragonfly give wing motor centers accurate population vector of prey direction. 2013. Proc. Natl. Acad. Sci. U.S.A. 110 p696–701.

8. Mischiati, M., Lin, HT., Herold, P., Imler, E., Olberg, R., and Leonardo, A. Internal models direct dragonfly interception steering. 2015. Nature 15, 333–8

9. Nordström, K., Douglas M. Bolzon, DM., and O’Carroll, DC. Spatial facilitation by a high-performance dragonfly target-detecting neuron. 2011. Biol. Lett. 7, 588–592.

10. O’Carroll, DC. Feature detecting neurons in dragonflie 1993. Nature 362, 541–543.

11. Olberg, RM. Identified target-selective visual interneurons descending from the dragonfly brain. 1986. J. Comp. Physiol. A, 159 p827–840.

12. Rieke, F., Warland, D., De Ruyter van Steveninck, R., and Bialek, W. Spikes: Exploring the neural code. 1997. MIT press, Cambridge.

13. Supple, JA., Pinto-Benito, D., Khoo, C., Wardill, TJ., Fabian, ST., Liu, M., Pusdekar, S., Galeano, D., Pan, J., Jiang, S., Wang, Y., Liu, L., Peng, H., Olberg, RM., and Gonzalez-Bellido, PT. Binocular Encoding in the Damselfly Pre-motor Target Tracking System. 2020. Curr. Biol. 30, p645–656.

14. Warzecha, AK., and Egelhaaf, M. Response latency of a motion-sensitive neuron in the fly visual system: dependence on stimulus parameters and physiological conditions. 2000. Vis. Res. 40, p2973–83

15. Wiederman, SD., Fabian, JM., Dunbier, JR. and O’Carroll, DC. A predictive focus of gain modulation encodes target trajectories in insect vision. 2017. eLife 6, e26478.

